# The response of a metapopulation to a changing environment

**DOI:** 10.1101/2021.09.17.460820

**Authors:** Nick Barton, Oluwafunmilola Olusanya

## Abstract

A species distributed across diverse environments may adapt to local conditions. We ask how quickly such a species changes its range in response to changed conditions. Szep et al (Evolution, 2021) used the infinite island model to find the stationary distribution of allele frequencies and deme sizes. We extend this to find how a metapopulation responds to changes in carrying capacity, selection strength, or migration rate when deme sizes are fixed. We further develop a “fixed-state” approximation. Under this approximation, polymorphism is only possible for a narrow range of habitat proportions when selection is weak compared to drift, but for a much wider range otherwise. When rates of selection or migration relative to drift change in a single deme of the metapopulation, the population takes a time of order 1/m to reach the new equilibrium. However, even with many loci, there can be substantial fluctuations in net adaptation, because at each locus, alleles randomly get lost or fixed. Thus, in a finite metapopulation, variation may gradually be lost by chance, even if it would persist in an infinite metapopulation. When conditions change across the whole metapopulation, there can be rapid change, which is predicted well by the fixed-state approximation.

## Introduction

Species must adapt to varied environments, whilst drawing on a common pool of genetic variation. Thus, there is a tension between selection that favours different alleles in different places, and the maintenance of diversity across the whole species. Local populations can only sustain themselves if they are sufficiently well-adapted; conversely, adaptation to conditions beyond the current niche can extend the range of the species.

These issues, which lie at the interface between ecology and evolution, have only quite recently attracted sustained theoretical attention. This ranges from studies of “evolutionary rescue”, typically of a single isolated deme [5, 6, 21], through to analyses of limits to a species’ range in one or two spatial dimensions [9, 12, 17]. Here, we consider an idealised metapopulation; in this island model, there is no explicit spatial structure. Nevertheless, we can ask whether the species’ range can extend over a variety of habitats, and examine how it responds dynamically to changing conditions – either in a single deme, or across the whole metapopulation.

We examine a simple model, in which directional selection favours alternative alleles in two different habitats. Provided that selection is stronger than migration, these alternative adaptations can be maintained despite gene flow. There is substantial literature on how heterogeneous selection can maintain diversity, beginning with Levene (1953). However, this is largely deterministic, neglecting random drift in small local populations. Here, we are primarily concerned with the erosion of adaptation by random drift within local demes - which can cause a substantial “drift load” even when the whole metapopulation is very large.

This paper is an extension of [19], which analysed the joint evolution of allele frequencies and deme sizes, in an island model with explicit density-dependent regulation; a diffusion approximation gave explicit formulae for the stationary distribution of an infinite metapopulation. Here, we extend this treatment to consider the evolution of individual demes, and of the whole metapopulation, as conditions change (directly through changes in selection and gene flow, and indirectly through their effect on the population size); we also consider fluctuations in a metapopulation with a limited number of demes, where variation can be lost by chance. We simplify the problem by assuming that deme sizes are fixed, independent of adaptedness (“soft selection”), but believe that the methods we introduce can be extended to allow density regulation (“hard selection”).

In principle, we can calculate the joint distribution of deme size and allele frequencies under the diffusion approximation. However, this is numerically challenging, since it involves a high dimensional partial differential equation; in any case, it can only be done for an infinite metapopulation, where the mean population size and allele frequencies across the population as a whole change deterministically, even though population sizes and allele frequencies within any deme follow a distribution. In order to go beyond mere simulation, we use the approximation that loci are typically near fixation; this is accurate if the number of non-native alleles that enter per generation is small. It allows us to follow the distribution of states of a finite metapopulation through time, which depends only on the rates of substitutions in either direction. This “fixed-state” approximation is an extension of models of “adaptive walks” (e.g. Orr [14], Trubenova et al [20]) to structured populations.

We first consider an infinite metapopulation, and determine the accuracy of the fixed-state approximation. We then apply the approximation to calculate the dynamics of a finite metapopulation, and to find how its equilibria depend on the number of demes. (In order for a non-trivial equilibrium to exist, we must allow a low rate of mutation to maintain variation in the long term). Finally, we show how metapopulations respond to changing conditions, focusing on changes that take the system between qualitatively different regimes.

## Model and Methods

We simulate a haploid population, assuming linkage equilibrium. Provided that selection is weak, this is accurate, and allows us to efficiently simulate large numbers of loci and demes; Szep et al (2021, SI C) examined the effects of linkage disequilibrium in this model, using individual-based simulations. We obtain analytical results by taking the diffusion limit, which also assumes weak selection, and then approximate this by assuming that demes are near fixation, which applies when there are few migrants (*Nm*<1). As is traditional in population genetics, we take the fundamental model to be the diffusion, since this captures the behaviour of a variety of particular life histories, and identifies the key dimensionless parameters.

### Simulations

Our baseline island model assumes that demes each have carrying capacity of *N* haploid individuals, and contribute equally to the migrant pool. A deme of size *N* is expected to lose a fraction *m* of individuals by emigration, and receives a Poisson distributed number of migrants, *Nm*\*, with expectation *Nm*. There are *L* biallelic loci, with the two alternative alleles labelled *X_i,k_* = 0 or 1; *i* labels the deme, and *k* the locus. Deme *i* is described by {*j*_*i*,1_,*j*_*i*,2_,…,*j*_*i,L*_}, where 0≤*j_i,k_*≤*N* is the number of copies of the ‘1’ allele at the *k*’th locus. That allele is favoured by selection *s_i_*, which we assume to be the same across loci; the marginal relative fitnesses are 1:*e^s_i_^*, and fitnesses multiply across loci. Under soft selection, loci evolve independently, and so it would be straightforward to extend to allow variation in selection across loci.

We assume linkage equilibrium (LE), and apply the Wright-Fisher model to each locus independently. After selection, allele frequencies are 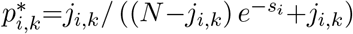, and after migration, 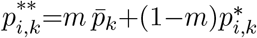 where 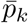 is the frequency averaged across all demes of the metapopulation. The new population in deme *i* consists of *N* individuals, the number of allele copies at locus *k* being binomially sampled with frequency 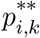. This procedure is accurate provided that s is not too large (<0.2, say), so that recombination shuffles genes faster than selection, drift, or migration build up associations between them (Szep et al., 2021, SI C).

A Mathematica notebook containing the simulation code and result is provided as a supplementary material (SM).

### Diffusion approximation

The diffusion approximation to this model describes the evolution of the joint distribution of allele frequencies across different demes, conditional on the mean allele frequency across the metapopulation [2]. A single deme follows a stochastic path governed by this distribution, whilst an infinite metapopulation represents the whole distribution, which evolves deterministically at the level of the metapopulation. The diffusion depends only on scaled parameters *Ns, Nm*.

Wright [22, 23] gave an explicit solution for the stationary distribution of allele frequencies:

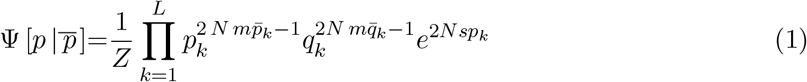

where *Z* is a normalising constant and *q*=1–*p*. Under this simple model of directional selection, allele frequencies evolve independently across demes and across loci, conditional on the mean allele frequencies, 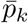. Equation (1) applies to a single deme; the subscript *i* was dropped for clarity. All demes that share the same parameters will follow the same distribution, in a given habitat, and so we can integrate over the distribution, and sum over habitats, to find the mean 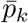. This allows us to solve fully for the stationary state.

### Fixed-state approximation

If the number of incoming alleles is small (*Nm*≪1) then the distribution of allele frequencies will be sharply peaked around 0 and 1. To a good approximation, populations are near fixation for one or other allele, and their state is determined by the rates of substitution in either direction. Since we will later be considering the stationary state of a finite metapopulation, we must include mutation, which we assume to be symmetric at rate *μ*. Then, the rate at which demes currently fixed for allele 0 substitute allele 1, λ_0_→_1_ (or vice versa, λ_1_→_0_) is the product of the number of ‘1’ (or ‘0’) alleles entering the population, and their individual fixation probability (see SM for details). Thus:

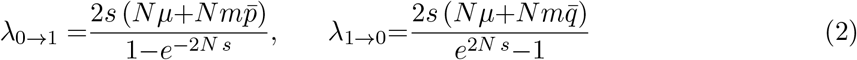

Different loci evolve independently, conditional on the numbers of migrants coming into the deme 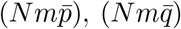. Note that when *Ns* and *Nm* are small, these rates are proportional to *m* in the neutral limit.

For an infinite metapopulation, and two habitats with selection *s*_1_,*s*_2_, with deme sizes fixed at *N* (i.e., soft selection), we can just follow the proportion of demes fixed for the ‘1’ allele in each habitat, *P*_1_, *P*_2_. Neglecting mutation, the rates given by eq. (2) lead immediately to:

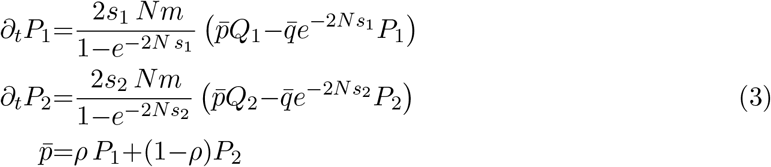

The first two equations involve the difference in net rates of substitution in each direction, where Q, P are the fraction of loci near fixation for 0, 1; 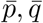 are the fraction of migrants with allele 1 vs 0, which can contribute to a substitution; and the fixation probabilities in each direction are in the ratio 1:*e*^-2*Ns*_1_^. Finally, the mean allele frequencies are a weighted average across habitats, which are in the proportions *ρ*:1-*ρ*.

These equations apply separately to each locus, but for simplicity, in numerical examples we will assume symmetric initial conditions, so that the proportion of demes fixed for the 1 allele in each habitat, *P*_1_, *P*_2_ are the same for all loci, and correspond to the proportion of loci fixed for the ‘1’ allele in each deme.

If the ‘1’ allele is favoured in habitat 1, but disfavoured in habitat 2 (i.e. *s*_2_<0<*s*_1_), and if neither habitat is too rare, then polymorphism is possible, with equilibrium frequency given by:

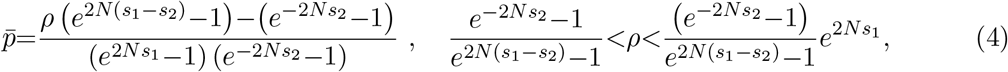

(as derived from eq. (2); see SM). If selection is weak relative to drift, polymorphism is possible only for a very narrow range of habitat proportions (see left of fig. A1 in Appendix A), whereas if it is strong, polymorphism is possible over a wide range (right of fig. A1 in Appendix A).

Suppose now that there are a finite number of demes, with *d_i_* having habitat *i*. At any one locus, the state of the metapopulation is described by the number of demes fixed for the ‘1’ allele, 0≤*k_i_*≤*d_i_*. For example, with two habitats, there are (*d*_1_+1) (*d*_2_+1) possible values for the state {*k*_1_,*k*_2_}. The probability of transitions between these states depends on the mean allele frequency across the metapopulation. With soft selection, where all demes have the same size *N*, this mean is just 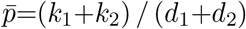. We can therefore calculate the transition matrix that governs the stochastic evolution of the metapopulation; the stationary state is given by the leading eigenvector of this matrix. With soft selection, each locus evolves independently, governed by this matrix, and so we can easily calculate the stochastic evolution of the metapopulation.

In Appendix B, we examine the accuracy of the fixed-state approximation under soft selection. This approximation applies in the limit of low migration, and identifies the failure of adaptation due to random drift.

## Results

### Evolution of a single deme

Consider a metapopulation, where *Nm* is small enough that populations are near fixation. If *Ns*_1_=1 in a rare habitat, represented in *ρ*=0.2 of the demes, and *Ns*_2_=-2 in the common habitat, then polymorphism will be maintained with 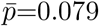 overall (eq. (4)). We begin by considering how a single deme responds to changes in its local conditions, for fixed 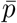, and so in fact, all that matters is the value of 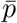. In the focal deme, allele frequencies will be in the ratio 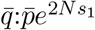 when *Nm*≪1, since that is the ratio of substitution rates in either direction; hence, the expected allele frequency in the rare habitat is 0.386 (lhs of fig. 1a). As *Nm* increases, the expected allele frequency decreases, approaching 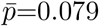 (rhs of fig. 1a). For given *Nm*, the expected allele frequency in the focal deme increases with *Ns*_1_ from 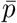 to 1, as selection becomes more effective (fig. 1b).

**Figure 1:**
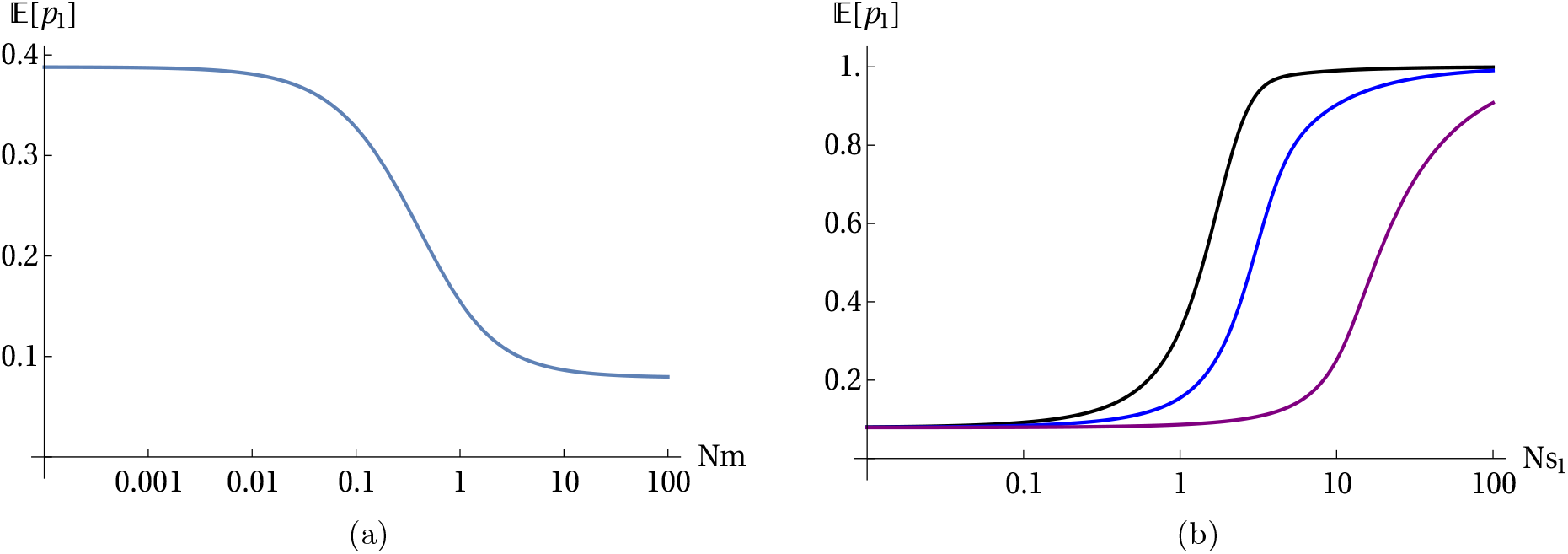
(a). Expected allele frequency vs *Nm* with 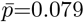, *Ns*_1_=1. (b). Expected allele frequency vs. *Ns*, for *Nm*=0.1, 1, 10 (black, blue, purple).

Figure 2a shows how the distribution of allele frequencies changes as *Nm* changes. If all loci start close to the frequency in the gene pool 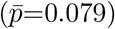 then with a low migration rate (*Nm*=0.05), even weak selection (*Ns*=1) can raise the mean substantially, to 0.355. However, this increase is slow, taking ~5000 generations, because it occurs through occasional substitutions, at a rate proportional to *m*=5×10^-4^ (eq. (3)). The population does mostly flip between fixation of one or other allele, giving a U-shaped frequency distribution (e.g. grey trajectory in fig. 2), and so the fixed-state approximation is quite close to the exact mean (orange vs. red at left). However, the average across even 100 loci fluctuates substantially (blue), implying that population fitness will fluctuate randomly, even when adaptation is highly polygenic.

**Figure 2:**
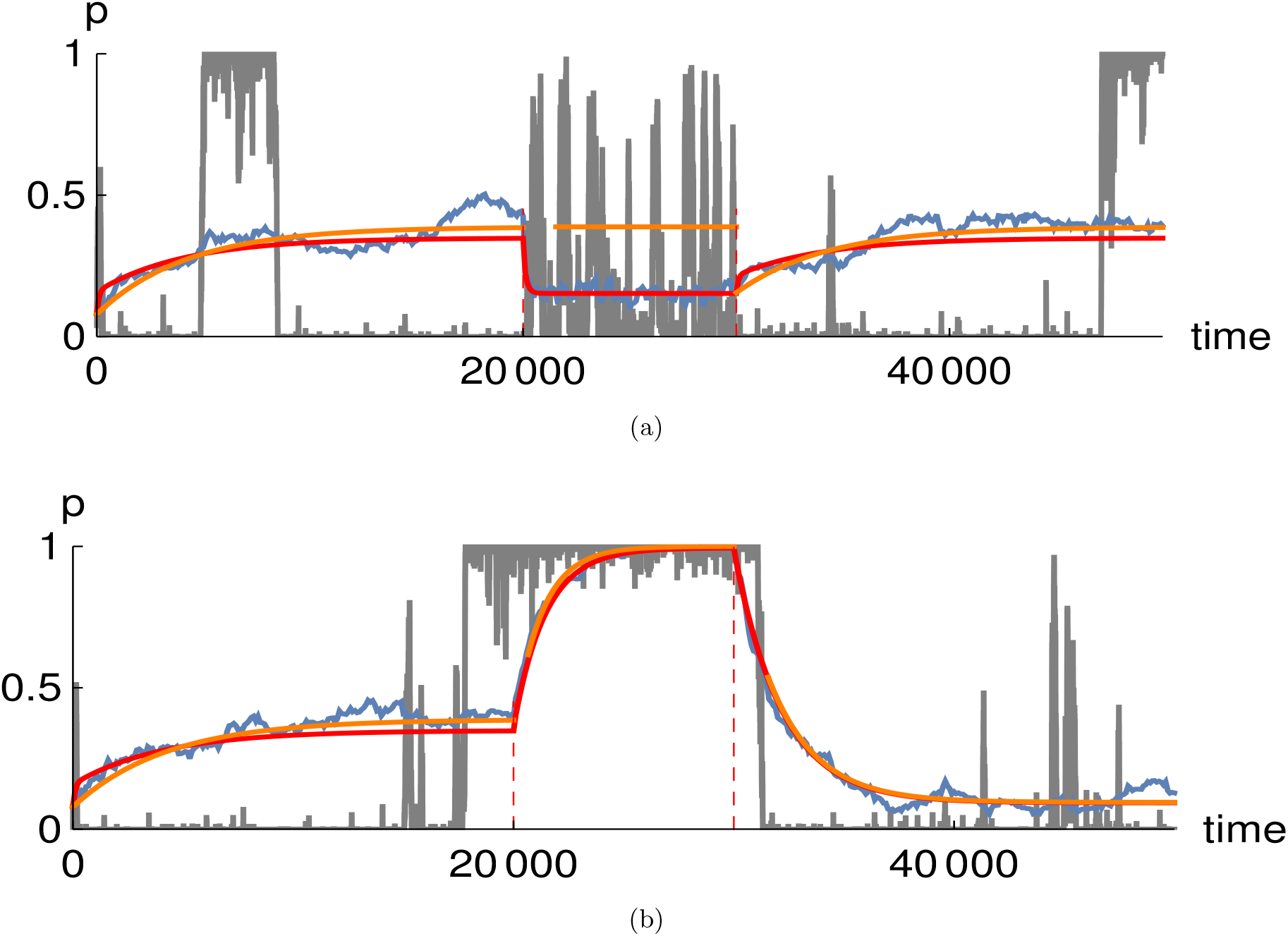
(a) Evolution of a single deme as *Nm* changes; *Ns*=1, 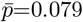, *L*=100 loci. Initially, *Nm*=0.05, and all loci are at 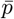. After 20, 000 generations, *Nm* increases to 1, and after another 10, 000 generations, it returns to *Nm*=0.05. The grey line shows allele frequency at a single locus, and the blue line shows the average over 100 loci. The red curve is the mean of the probability distribution, calculated exactly using the Wright-Fisher transition matrix. The orange curve is the fixed-state approximation (eq. (3)), which is accurate only for *Nm*≪1. (b) The same as (a), but for *Ns* changing from 1 to 10 at 20, 000 generations, and then to 0.1 at 30, 000 generations; *Nm*=0.05 throughout.

At 20,000 generations, the number of migrants increases to *Nm*=1, and the mean allele frequency is quickly pulled down towards that in the gene pool, to 0.155. The fixed-state approximation is the limit of low migration, and so is independent of *Nm* (see fig. A2). Indeed, allele frequencies are now often intermediate, and so this approximation fails (orange vs. red, fig. 2, middle). Nevertheless, it does give the important intuition that rates of change are proportional to migration, which is now *m*=0.01, implying a 1/*m*~100 generation timescale for response of the population mean. In this model, variance is maintained by migration, and so the response to selection is proportional to *m*: we can see this in eq. (3), where rates of change are proportional to *m*, for given *Ns*. After *Nm* returns to the original low value at 30,000 generations, there is a slow return to the original bimodal distribution, again captured by the fixed-state approximation (orange vs red at right of fig. 2).

Figure 2b shows the response to changes in *Ns*, which could arise through changes in selection strength, and/or changes in local effective population size. In this example, *Nm*=0.05 throughout, and so the fixed-state approximation is accurate (orange vs red curves). The timescale is again set by *m*, which determines the rate at which variation is introduced into local demes. Since *m*=5×10^-4^, it takes thousands of generations for the proportion of loci fixed for the ‘1’ allele to respond to changes in selection strength.

Figure 3 shows the time taken for a population to respond to changes in *Nm* (fig. 3a) or *Ns* (fig. 3b), as a function of the other parameter. In each figure, the two curves show the time to respond to changes in either direction. As we saw in fig. 2a, an increase in *Nm* (lower plot of fig. 3a) causes a much faster response than a decrease (upper plot of fig. 3a), simply because high gene flow introduces more genetic variance. However, if selection is very strong, the response time becomes similar in either direction, and decreases in proportion to *Ns* (right of fig. 3a). The response to changes in *Ns* take somewhat longer for an increase than a decrease (compare upper vs lower plot of fig. 3b), but the main pattern here is that the response time decreases in proportion to *Nm*.

**Figure 3:**
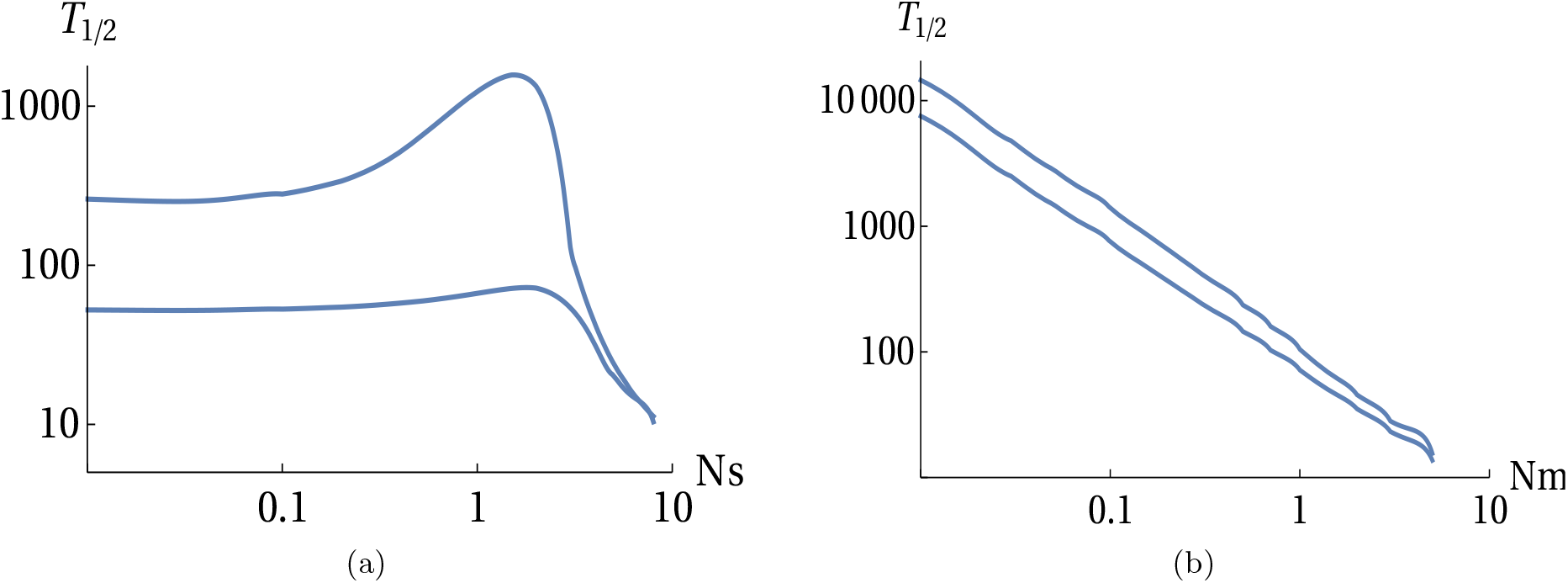
The time to make half of the response to a change in parameters. For both plots, 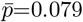. Values were calculated using a transition matrix with *N*=100. Details of the calculation are in the SM. (a) *Nm* shifts from 0.05 to 1 or from 1 to 0.05 (lower, upper curves, resp.), for varying *Ns*. (b) *Ns* shifts from 0.1 to 1 or from 1 to 0.1 (upper, lower curves, resp.) for varying *Nm*.

### Evolution of a metapopulation

We begin by considering the stationary state of a metapopulation, extending Szep et al [19] by allowing a finite number of demes - in which case, a low rate of mutation is required to maintain variation in the long term. We then give an example that shows how variation is lost, as loci fix across the whole metapopulation. Finally, we give examples (analogous to fig. 2), showing the response when parameters change across the whole metapopulation.

### Stationary state of a finite metapopulation in the limit of small *Nm*

Szep et al (2021, Fig. 2) show that with soft selection, polymorphism can be maintained in an infinite metapopulation, provided that selection is sufficiently strong. With symmetric selection (*s*_1_=*s*_2_), this requires 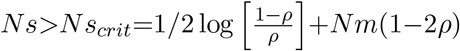; the first term is derived from the fixed-state approximation, in the limit *Nm*≪1, and the second from the deterministic model, which requires *s*>*m*(1-2*ρ*) for polymorphism (see Szep et al 2021, immediately above “Hard selection”). In a metapopulation with a finite number of demes, variation must ultimately be lost: we must include mutation to allow a non-trivial stationary state. In this section, we examine how the outcome depends on the relative rates of selection and drift (*Ns*) and on the relative rates of mutation and migration (*μ/m*). In particular, we show that with sufficiently many demes, the outcome is insensitive to the mutation rate.

Figure 4 shows the stationary state in the limit of small *Nm*, derived using the fixed-state approximation. The top row of fig. 4 shows how the fraction of demes fixed at equilibrium in a rare and common habitat (i.e. 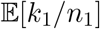 and 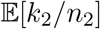 respectively) depend on the strength of selection when mutation is appreciable (fig. 4a) compared to when it is weak (fig. 4b), where *k_i_*, *i*=1,2, are as defined earlier, *n*_1_ and *n*_2_ are the total number of demes in the rare and common habitats respectively, *n*=*n*_1_+*n*_2_ is the total number of demes in the metapopulation and mutation is assumed symmetric. Out of all demes, *n*, in the metapopulation, the focal allele is favoured in 20% of demes (constituting the rare habitat) so that *n*_1_ = 0.2 *n*, and disfavored twice as strongly in the remaining 80% of demes (constituting the common habitat) so that *n*_2_ = 0.8 *n*. The blue (red) color represents dynamics in the rare (common) habitat when we have a finite metapopulation (i.e. *n* = 50, 100, 200, and 400 demes represented by the solid, dashed, dotted and dot-dashed lines respectively). Whereas, the purple (orange) color represent dynamics in the rare (common) habitat when we have an infinite metapopulation.

**Figure 4:**
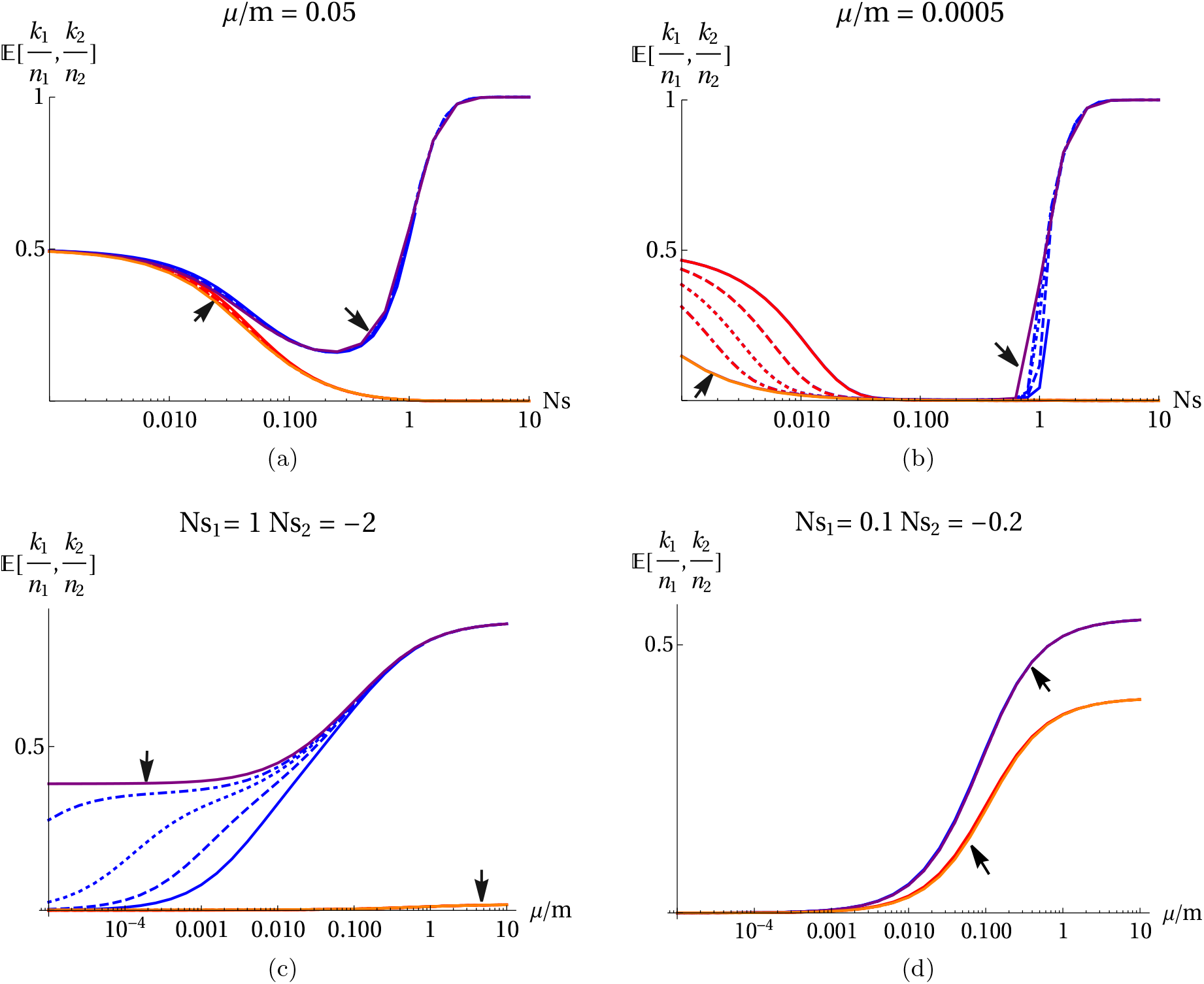
The fraction of demes in each of the two habitats (rare habitat – blue, common habitat – red) that are fixed, as a function of selection strength (*Ns*, top row) and the rate of mutation, relative to migration (*μ/m*, bottom row). The focal allele is favoured by selection *Ns*_1_ in 20% of demes (i.e. in the rare habitat – blue), and disfavoured by selection *Ns*_2_ =–2*Ns*_1_ in the remaining 80% of demes (i.e. in the common habitat – red). In each plot, equilibria for 50, 100, 200 and 400 demes are superimposed (solid, dashed, dotted and dot-dashed lines), together with the limit for an infinite metapopulation (solid purple and orange lines indicated by arrows).

When mutation is appreciable (*μ/m* = 0.05, fig. 4a), the focal allele is unlikely to be lost by chance (blue colors). Furthermore, the equilibrium is insensitive to the number of demes, and close to the solution for an infinite population (as can be seen from the indistinguishability of the various lines for different *N*). When selection is strong (right of fig. 4a, all demes are fixed for the favoured allele, whereas when selection is negligible, on average half of the demes are fixed for each allele. In-between (i.e. 0.1<*Ns*<1), the allele favoured in the rare habitat becomes rare, being pulled to low frequency by migration from the commoner habitat, where it is more strongly disfavoured. When mutation is weak relative to migration (as is likely in nature, i.e. *μ/m*=0.0005; fig. 4b), this pattern is exaggerated. Above a critical value, 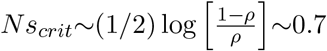, polymorphism can be maintained by divergent selection, despite drift and gene flow. The equilibrium for an infinite population (purple solid line, indicated by arrow) gives an upper bound, but stochastic loss from a finite set of demes reduces the expected frequency, and increases the critical *Ns_crit_* (solid, dashed, dotted and dot-dashed blue lines around *Ns*~1, for 50,100,200 and 400 demes respectively). There is a wide region (0.03<*Ns*<0.7) where the allele is almost absent, being swamped by gene flow. However, for very weak selection (fig. 4b, lhs), the frequency of the allele increases towards the symmetric neutral equilibrium i.e. 0.5. In this regime, the frequencies in the two habitats are almost identical, and cannot be distinguished in the figure. Furthermore, in this regime (left of fig. 4a), although selection is negligible within demes (*Ns*<0.1), migration is much faster than mutation, and so selection over the whole metapopulation is effective in eliminating the allele that is deleterious in most demes. Therefore in this scenario where mutation is rarer than migration, and selection is weak relative to drift within a single deme (i.e. *μ*≪*m*, *Ns*<0.1), selection is nevertheless still effective at the level of the whole metapopulation and is especially so in the habitat which has more demes (left of fig. 4a).

The bottom row of fig. 4 shows the dependence on the relative rates of mutation versus migration, *μ/m*. With high mutation rates, the equilibrium approaches a fraction 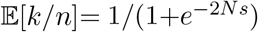, given by the ratio of fixation probabilities (eq. (2)) in the fixed-state approximation. There is strong divergence when *Ns*_1_=1 (right of fig. 4c), and weaker divergence when selection is weak (fig. 4d, *Ns*_1_=0.1). With moderately strong selection (fig. 4c), the allele that is less favoured overall is lost from the common habitat, independent of the number of demes and mutation rate (orange line with arrow head). In the rare habitat, with weak mutation (left of fig. 4c), the locally favoured allele can be fixed in nearly half the demes in an infinite metapopulation (purple solid line with arrow head), but tends to be lost by chance from finite metapopulations, even with several hundred demes (solid, dashed and dotted blue lines). When selection is weak relative to local deme size (fig. 4d), selection can still be effective over the whole metapopulation, eliminating the allele that is disfavoured overall (left of fig. 4d). However, when mutation becomes comparable with migration, polymorphism is maintained by mutation pressure, with some bias between habitats caused by weak selection (right of fig. 4d).

We focus on the regime where selection is comparable to drift (*Ns*_1_~1), and mutation is weak. This corresponds to the right half of fig. 4b (0.1<*Ns*_1_), and the middle of fig. 4c (10^-4^<*μ/m*<0.1). Then, as long as mutation is not extremely small, and there are enough demes, the stationary state is close to that in an infinite metapopulation (compare blue dashed and dotted lines with purple line in fig. 4c). However, note that with weak mutation (*μ/m* ~ 10^-^-10^-3^, say), the locally favoured allele tends to be lost even when there are several hundred demes.

### Loss of diversity in a finite metapopulation

When deme sizes are fixed, and numbers of migrants are low enough that loci are typically fixed for one or other allele, the state of the metapopulation at each locus can be described by the number of demes, *k_i_*, in each habitat, *i*, that are fixed for the ‘1’ allele. The distribution of *k_i_* evolves according to a transition matrix, and each locus follows an independent realisation of the same stochastic process. Figure A3 and A4 in Appendix C compares the dynamics of the fixed-state approximation with simulations, to illustrate its accuracy. For *Nm* = 0.05 (fig. A3), there is reasonable agreement between simulations and the fixed state approximation and for lower *Nm* (i.e. *Nm* = 0.01), agreement is even more close (fig. A4). In both cases, variation is lost faster than predicted by the fixed-state approximation (compare red and black lines of fig. A3a and A4a), because migration tends to swamp adaptive divergence. The timescale is inversely proportional to *m*, and is therefore slow. Here, we are focusing on the slow loss of adaptation through random drift in small populations; with higher migration rates, swamping by gene flow causes additional, faster, degradation.

Note that because the number of demes is limited, and because each deme flips between fixation for alternative alleles, there is substantial variability in average allele frequency between loci (grey lines). Therefore, adaptation is lost slightly faster in a finite than in an infinite metapopulation (compare black and magenta lines in both fig. A3a and A3b, which both derive from the fixed-state approximation). Nevertheless, the overall mean, averaged over 40 loci, changes smoothly and predictably (red curves in fig. A3 and A4). We assume no mutation, and so all variation will inevitably be lost. However, because the total population is large (i.e. 100 demes × 50 individuals per deme = 5000 individuals), and because very low migration rate increases the effective size of the whole metapopulation, loss across the whole metapopulation is extremely slow: none of the 40 loci fix during the 10^4^ generations shown in fig. A3 and A4 (grey lines).

### Response to changing conditions across the metapopulation

Figure 5 shows some examples of the response to a change in conditions when parameters change gradually in all demes of the rare habitat. We focus on the rarer habitat since we are mostly concerned with adaptation in the rare habitat and its degradation by gene flow.

**Figure 5:**
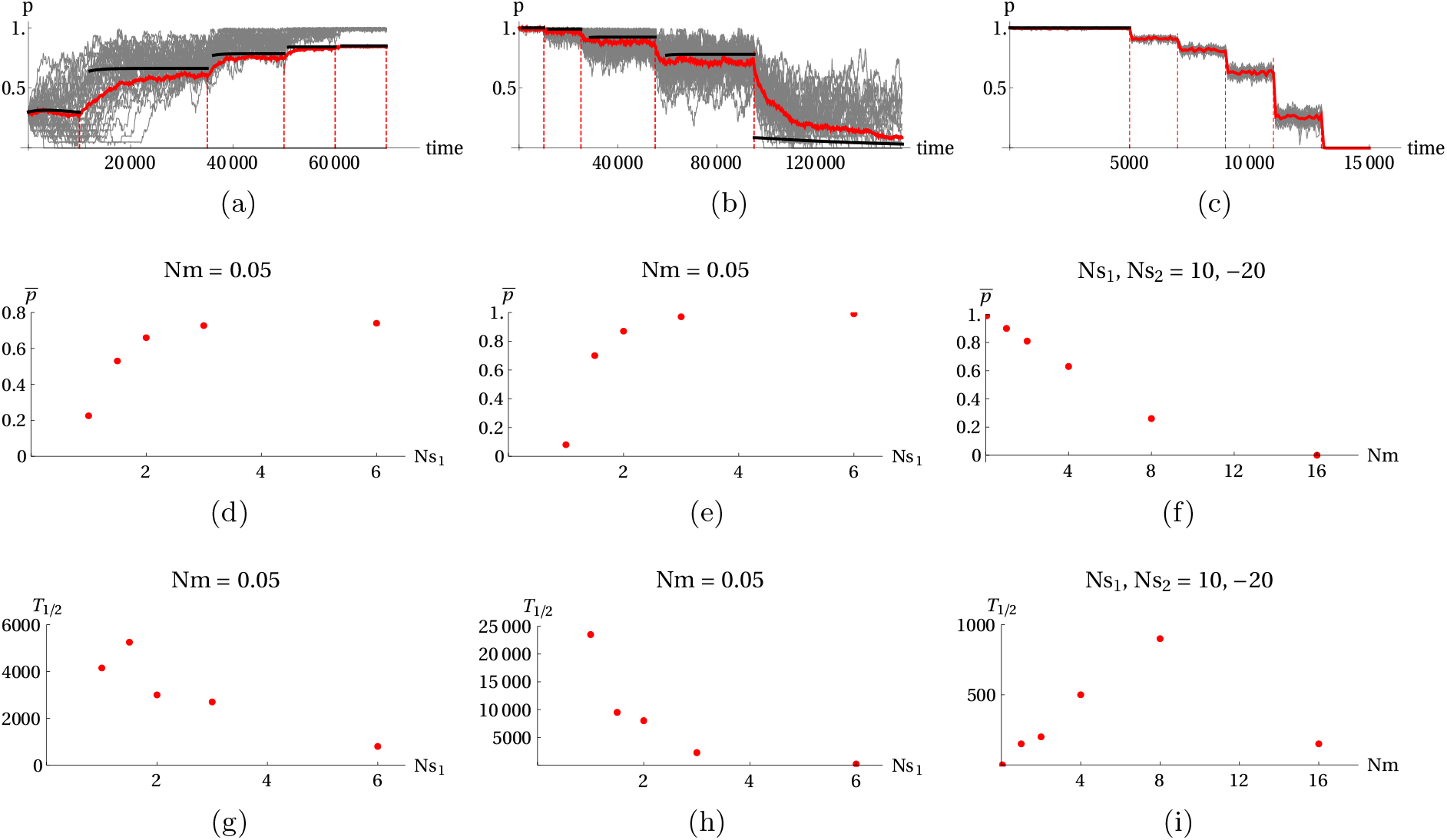
(A)-(C) Response of a metapopulation to changing conditions. Grey lines show the allele frequencies, averaged over the 20 demes in the rare habitat, at each 40 loci; the red lines show the overall mean in the rare habitat. The black line shows the prediction in the limit of small *Nm* (i.e. the fixed state approximation). In figure A. *Nm* is kept fixed at 0.05 and *Ns*_1_ is increased gradually from 1→1.5→2→3→6 with *Ns*_2_ fixed at 2. In figure B., we have the reverse of figure A. where *Nm* is again kept fixed at 0.05, *Ns*_2_ kept fixed at 2 and *Ns*_1_ is now decreased gradually from 6→3→2→1.5→1. In figure C. *Ns*_1_ and *Ns*_2_ are kept fixed (at 10 and –20 respectively) and *Nm* is gradually increased from 0.05→1→2→4→8→16. (D)-(F) Plot of 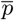 against the changing parameter. (G)-(I) Plot of *T*_1/2_ (i.e. half time to reach the new equilibrium 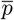) as a function of the changing parameter. Each point in fig. 5d – 5i is based on a single replicate. For all plots, simulations are run with 100 demes, *N* = 50, *L* = 40 loci and *ρ* = 0.2.

Figure 5a, 5d and 5g respectively show the consequence of a gradual increase in *Ns*_1_ on the distribution of allele frequencies in the rare habitat, the equilibrium allele frequency, 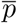 (averaged across all demes in the rare habitat) and the half time, *T*_1/2_, taken to reach this equilibrium mean frequency. We begin initially with selection comparable to drift (*Ns* = 1) and with a fraction of demes fixed for the locally favored allele, in the proportions predicted by an infinite metapopulation. After several thousand generations when the local allele has reached an equilibrium, we gradually increase *Ns*_1_ until a new equilibrium is obtained and do this continuously until all loci in the rare habitat reach (near) perfect adaptation (fig. 5a, grey lines). With selection comparable to drift, there is a substantial variation across loci with the 40 individual loci following markedly different paths and a chance loss of 6 of the loci (due to the effect of drift). As selection becomes stronger, the remaining polymorphic loci rise in frequency. Although there is still considerable variation in the rates of increase across loci (20,000 - 50,000 generations, fig. 5a), the overall mean, 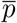 is approached quite smoothly (red lines). These new equilibrium mean values have a positive dependence on *Ns*_1_ (fig. 5d) and it takes a shorter time to reach a new equilibrium with stronger selection (fig. 5g).

Figure 5b, 5e and 5h correspondingly show a similar scenario as the above but with *Ns*_1_ now changing in the opposite direction (i.e. from high to low value). Initially, with selection much stronger than drift, all 40 loci are at a considerably high frequency (near 1) with less variability amongst them (lhs of fig. 5b, grey lines) so that the equilibrium mean allele frequency ~1 (fig. 5e, rhs). Also, because of this strong selection, the new equilibrium is approached rather fast (fig. 5h, rhs). As *Ns*_1_ declines, there is an apparent increase in the variability amongst loci so that with *Ns*_1_ = 1 drift becomes sufficient enough to cause the loss of three quarters of the loci (rhs of fig. 5b, grey lines). Because only 10/40 of the loci remain polymorphic, the overall mean is ~0.1 (fig. 5e, lhs).

fig. 5c, 5f and 5i show the response (of the distribution of allele frequency, the equilibrium mean frequency, 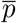 and the half time to 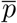) to an increase in *Nm* with *Ns*_1_, *Ns*_2_ = 10, −20 throughout. Since selection is generally strong relative to drift (*Ns*≫1) and gene flow is initially low (*Nm* = 0.05, fig. 5c, lhs), the rare habitat is initially perfectly adapted with 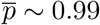 (fig. 5f, lhs). However, as *Nm* increases, gene flow gradually degrades adaptation as the focal allele is rapidly swamped until it is lost from the population after ~ 13,000 generations (fig. 5c, rhs). Since 0 loci remain polymorphic at this point, 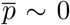 (fig. 5f, rhs). The half time to reach the new equilibrium values depend non-monotonically on *Nm*.

In fig. 5a – 5c, the dynamics are closely predicted by the small *Nm* limit (compare red and black curves) which is based simply on the rates of substitution in both direction.

## Discussion

Our analysis uses simulation, the diffusion approximation and the “fixed-state” approximation to understand how drift degrades adaptation in a finite metapopulation as well as how a finite metapopulation changes through time, as it responds to changes in both local and global conditions. The “fixed state” approximation applies either where variation is due to mutation (when it is plausible that *Nμ*<1 within local demes, or even at the level of the whole population), or when variation is maintained by divergent selection across the whole metapopulation, but migration is low relative to drift (*Nm*<1).

Using the fixed-state approximation, it was shown that when selection is weaker than drift (i.e. *Ns*<1), polymorphism can only be maintained for a very narrow range of habitat proportions (fig. A1). However with strong selection, this range becomes much wider. Also, using both simulations and the fixed state approximation, we showed that when conditions in a single deme of the metapopulation change, the population responds on a short time scale of order 1/*m*, simply because in the regime we study, local genetic variance is maintained by migration. Variation may be temporarily lost as local conditions change, but can quickly be recovered. On the other hand, when conditions change across the whole metapopulation, variation that was maintained by divergent selection can be permanently lost, and is only slowly recovered by mutation. Even under constant conditions, variation at a locus can be lost by chance, unless there are a very large number of demes.

To simplify our analysis, we assumed an island model, with a large number of spatially equivalent demes (i.e. soft selection). This is unlikely to be the case in nature, but may nevertheless capture the behaviour of spatially extended populations if there is long-range migration, which can introduce locally adaptive alleles from a distant habitat. It may be that a leptokurtic dispersal distribution can allow efficient adaptation, if locally favoured alleles are not swamped, and yet can be recovered by occasional long-range migration [1, 10].

Our analysis can however be further extended to hard selection, by including explicit density regulation; Szep et al [19] showed that one can still apply the diffusion approximation, provided that growth rates are not too high. With hard selection, substitution rates now depend on deme size through *Ns*, and through the number of immigrant alleles, 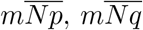. This dependence can be approximated by assuming that the population size is determined by the genetic load. Sachdeva et al [18] and Szep et al [19] refer to this as the “semi-deterministic” approximation which is accurate when demographic stochasticity is weak. One can apply the “fixed-state” approximation by further assuming that there are enough loci that the mean load is proportional to the mean across loci of the number of demes fixed for one or the other allele. The transition matrix can then be calculated as before, but is now a function of the population sizes in the two habitats, {*N*_1_,*N*_2_} which both depend on the current state via the load. The key assumption here is that with enough loci, the population sizes change almost deterministically, following the distribution of states across loci. One complication with hard selection however is the existence of multiple stable equilibria: changing conditions would not just cause equilibria to shift but also changes the rates of transitions between equilibria.

A key assumption in our analysis is that selection is directional: in a given environment, alleles experience a fixed selection pressure, which tends to drive out variation. More often, selection may favour an intermediate optimum for a quantitative trait (the so called stabilizing selection framework), such that when the mean is well-adapted, alleles are close to neutral. Our modelling framework can describe this case, but it is much more complex, since many different allele combinations can achieve the same optimum. However, if selection on each allele is weak (*Ns*<1), then the infinitesimal model [3] applies, and can also describe the population dynamics [4]. Local adaptation may be possible under higher migration rates in such a regime.

We have shown that it is very difficult for directional selection to maintain local adaptations when selection is weak, relative to deme size (*Ns*~1). Migration must also be weak if it is not to swamp adaptation, in which case alleles are typically near to fixation, limiting the genetic variance available for adaptation, and preventing the recovery of variation lost to random drift. This contrasts with *global* adaptation: selection can be effective across the whole metapopulation, even if selection is weaker than drift within local demes (*Ns*<1), provided that there is sufficient gene flow (*Nm*>1) [2]. Thus, we expect local adaptation to depend on relatively more strongly selected alleles than global adaptations [24].

In this work, we have also introduced a novel approach to understanding the dynamical evolution of metapopulations. Although the full behaviour requires simulation, the diffusion approximation allows the stationary state to be calculated, and identifies the key dimensionless parameters. Moreover, when migration is rare, we can use a fixed state approximation that connects population genetics with models of adaptive walks [14].

Our work suggests several open questions that invite theoretical study. First, although we show that local adaptation requires that directional selection be stronger than drift within demes, that may not be the case with stabilizing selection. Under the infinitesimal model [3], genetic variance may be due to weakly selected alleles (*Ns*<1), and yet can still sustain adaptation of polygenic traits. Second, local adaptation may be greatly impeded by hard selection: if maladapted populations collapse, they cannot be the site of future adaptation. Testing the theory in nature is challenging, because it requires measurement of fitness as well as genetic data. Nevertheless, it might be possible to find how manipulation of local deme size and gene flow (or natural variation in these parameters) alters fitness. If we know of loci responsible for local adaptation (e.g Jones et al [11], Pfeifer et al [16]), then the theory developed here can be applied more directly, though in practice it would need to be modified to account for actual spatial structure.

In conclusion, the methods introduced in this study allow us to explore how species adapt to diverse environments: we can find how organisms expand their range to a wider span of environments, through local adaptation that is sustained by variation maintained across the whole metapopulation. Many questions remain to be studied within this framework: for example, how populations adapt to large numbers of diverse habitats; whether population regulation (“hard selection”) leads to a feedback that impedes adaptation; and whether genetic variation can be better sustained under stabilizing selection towards a varying optimum, rather than the directional selection studied here. We believe that, despite the absence of explicit spatial structure, this approach will be a fruitful way to better understand what limits the range of environments that a species can occupy.

## Appendix

### A. Range of habitat proportions for which polymorphism is possible

**Figure A1:**
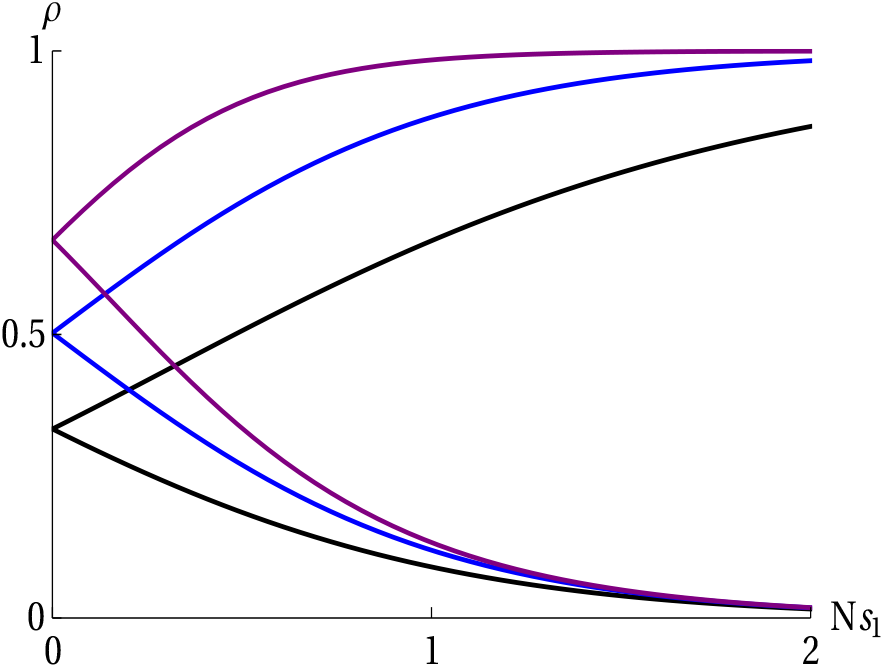
Bounds on the proportions of habitat 1, *ρ*, between which polymorphism is possible, as a function of the strength of selection in that habitat, *Ns*_1_. For a given *Ns*_1_, *ρ* has to lie between the two curves for polymorphism to be maintained. The three sets of bounds correspond to *Ns*_2_/*Ns*_1_ = 0.5,1, 2 (black, blue and purple respectively). These results apply in the limit of low migration, and soft selection.

### B. Accuracy of the fixed-state approximation

Here, we compare the the mean allele frequency in an infinite metapopulation under the diffusion approximation with the fixed-state approximation for different *Nm* values. As expected, the accuracy of the fixed state approximation holds only for small *Nm*.

**Figure A2:**
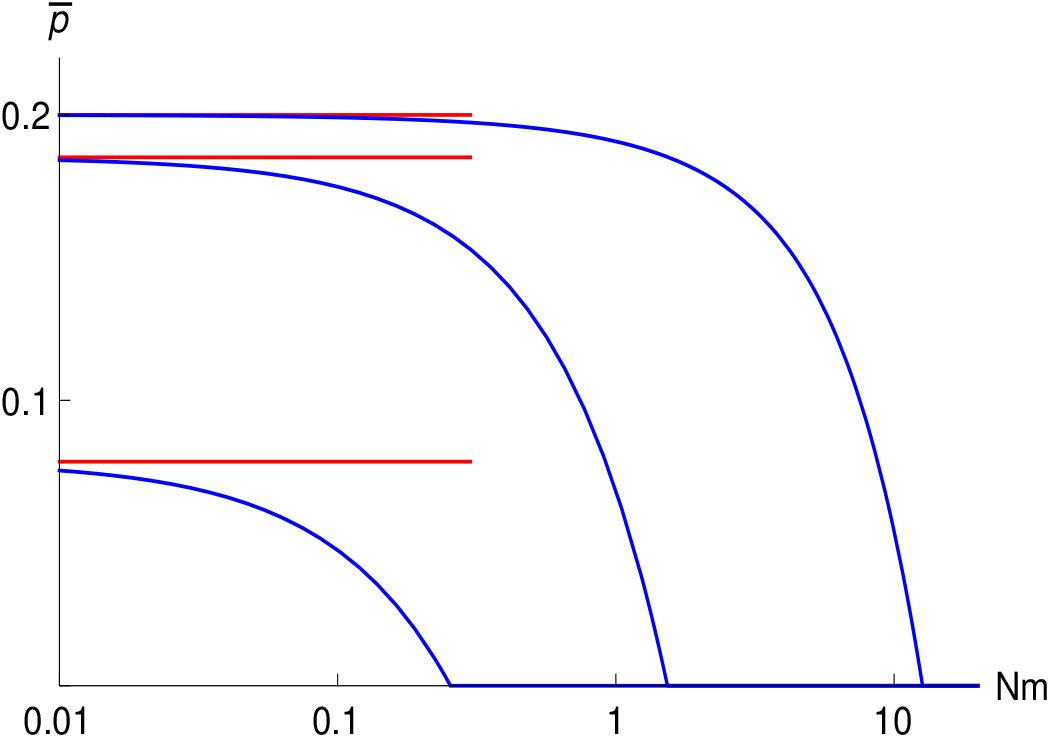
The mean allele frequency in an infinite metapopulation, plotted against *Nm; ρ* = 0.2, *Ns*_1_, *Ns*_2_ = 1, –2 (lower curve) 2, –4 (middle curve) or 10, −20 (upper curve). The fixed-state approximation, which applies for small *Nm*, is shown by the red lines.Loss of diversity in a finite population

### C. Loss of diversity in a finite population

**Figure A3:**
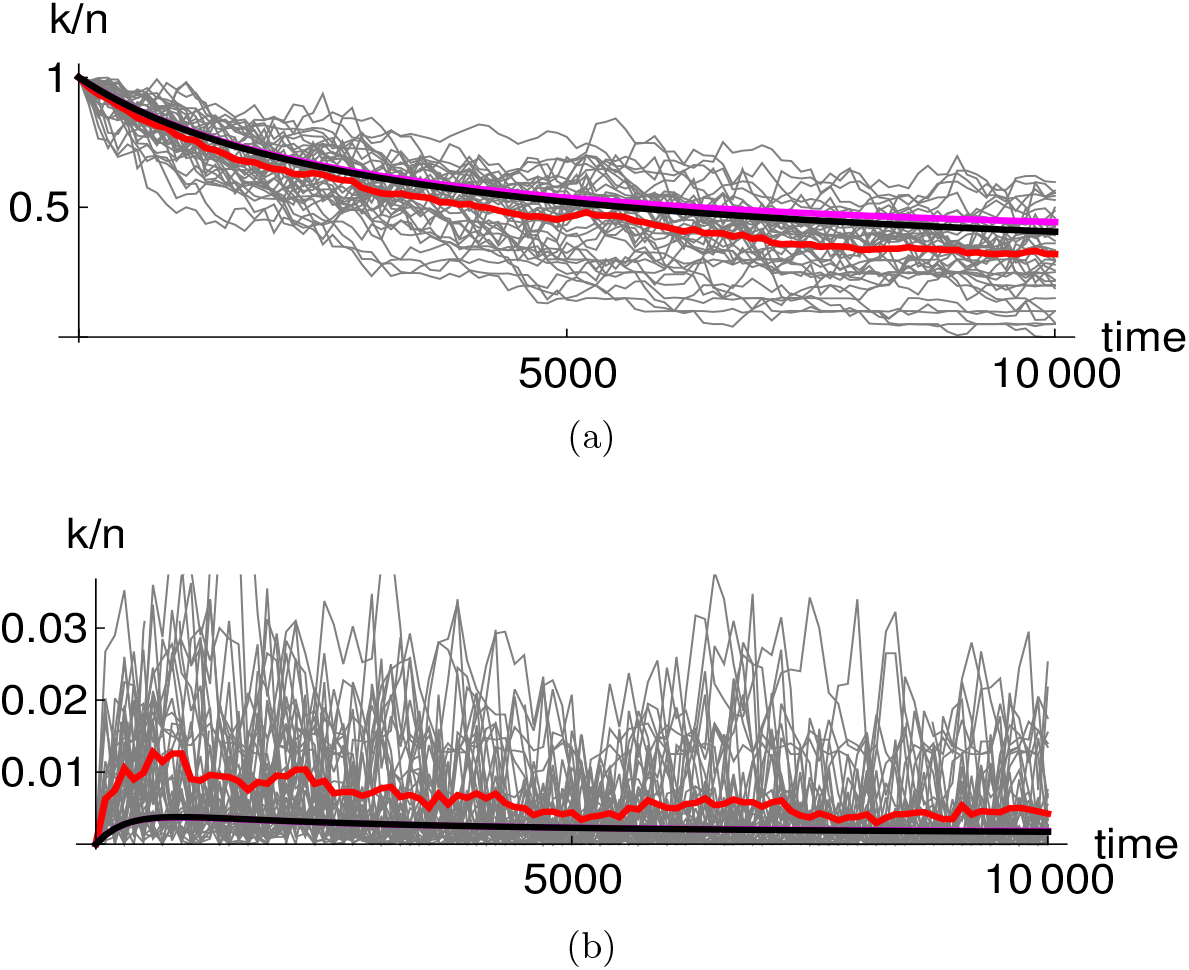
Loss of diversity in a metapopulation of 100 demes, which is initially perfectly adapted. (a) Mean allele frequency is plotted against time, in the 20 demes where the focal allele is favoured and, (b) in the 80 demes where it is not. Thin grey lines show allele frequencies at 40 loci, averaged over demes; the red line shows the overall mean. The black curve shows the fixed-state approximation, for a finite metapopulation, and the magenta line, for an infinite metapopulation. Simulations are for *N* = 50, *Nm* = 0.05, *s*_1,2_ = {0.02, –0.04}; thus, *Ns*_1,2_ = {1, –2}, so that selection and drift are of similar magnitude.

**Figure A4:**
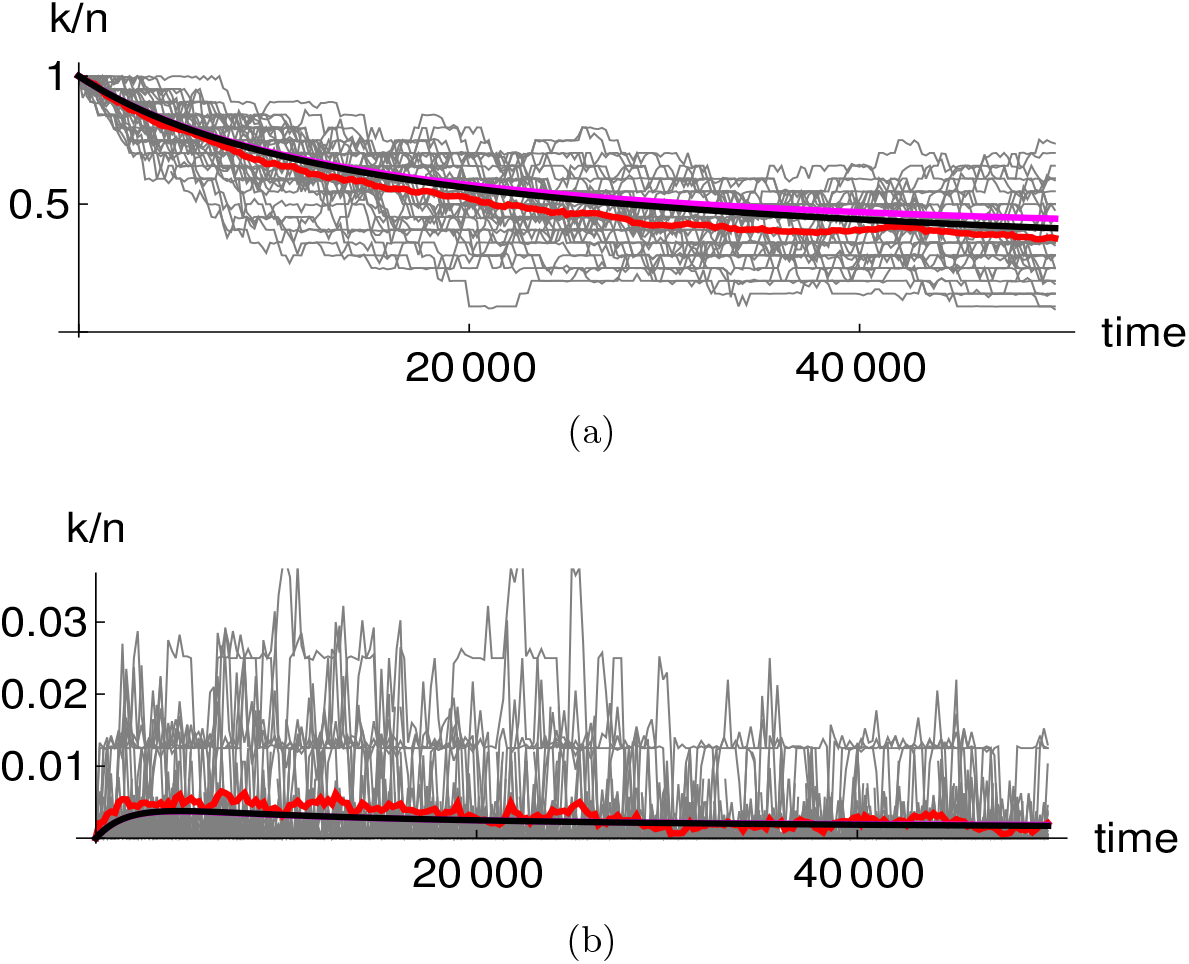
This is identical to fig. A3, except that *Nm*=0.01, and the timescale is correspondingly longer. The fixed-state approximation is more accurate with a lower number of migrants.

### D. Distribution of mean allele frequency

**Figure A5:**
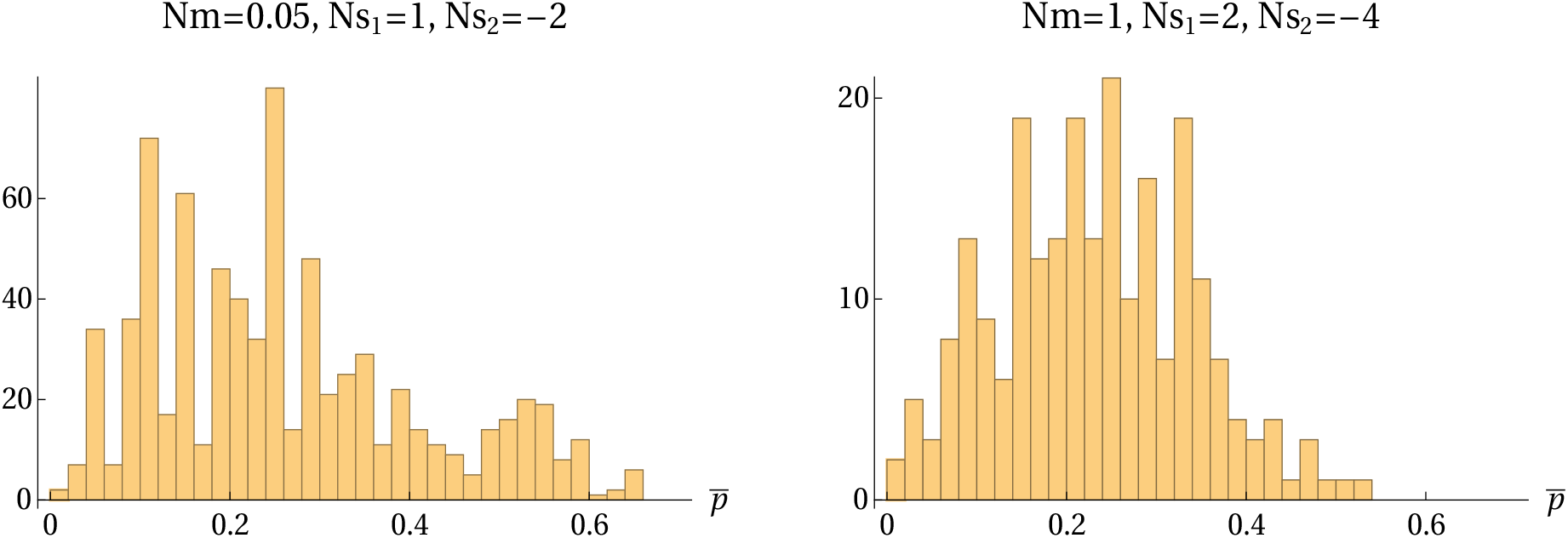
The distribution of allele frequencies, averaged over the 20 demes in the rare habitat, conditional on polymorphism, and accumulated over generations 8, 000 and 8, 100, to 10, 000.

